# Compensatory responses to glaucoma pathology in the dorsolateral geniculate nucleus

**DOI:** 10.1101/2025.09.23.678033

**Authors:** Shaylah McCool, Arnav Jain, Jennie C. Smith, Matthew J. Van Hook

**Affiliations:** Department of Ophthalmology & Visual Sciences, University of Nebraska Medical Center, Omaha NE 68198 USA; Department of Pharmacology & Experimental Neuroscience, University of Nebraska Medical Center, Omaha NE 68198 USA; Department of Cellular and Integrative Physiology, University of Nebraska Medical Center, Omaha NE 68198 USA

**Keywords:** Glaucoma, dorsolateral geniculate nucleus, plasticity, synapse, inhibition, excitability

## Abstract

Glaucoma disrupts the conveyance of retinal signals to visual regions of the brain such as the dorsolateral geniculate nucleus (dLGN) due to degeneration of retinal ganglion cells (RGCs) and their axons. Although plasticity during development allows altered visual experience to modulate dLGN synapses and excitability, evidence for experience-dependent dLGN plasticity in adults is limited. However, glaucoma might trigger compensatory plasticity in adult dLGN, thereby compensating for diminished RGC synaptic drive. Here, we tested this using aged (11-15 month-old) DBA/2J mice, which develop high intraocular pressure and glaucoma. In brain slice recordings, we found that diminished RGC inputs could drive robust action potential firing in dLGN relay neurons that was comparable to controls. This was accompanied by increased intrinsic excitability and decreased magnitude of sustained inhibitory currents from GABA spillover to extrasynaptic receptors. These results implicate multiple cellular and synaptic mechanisms that support signaling despite the diminished RGC inputs in glaucoma.

**HIGHLIGHTS:**

- Retinogeniculate synapse strength in the dLGN is diminished in the DBA/2J mouse model of glaucoma
- Despite a loss of synaptic strength, retinogeniculate synapses drive robust action potential firing in dLGN relay neurons, suggestive of homeostatic compensation
- dLGN relay neurons from DBA/2J mice have an increase in intrinsic excitability, supporting action potential generation
- Reduced extrasynaptic sustained inhibition in DBA/2J dLGN relay neurons can also support synaptically driven action potential firing

## INTRODUCTION

The dorsolateral geniculate nucleus (dLGN) of the thalamus is a critical waypoint for information traveling from the retina to the primary visual cortex (V1) for conscious vision ^1–3^. In the dLGN, retinal ganglion cells make strong, high release probability excitatory synapses onto thalamocortical (TC) relay neurons. TC neurons integrate those inputs along with feedforward inhibition from local interneurons, feedback inhibition from thalamic reticular nucleus, and excitation from Layer 6 of V1 to drive action potential output. Synaptic and neuronal function in the developing dLGN is subject to experience-dependent plasticity during a critical period occurring around postnatal days 20-35 in mice ^4,5^. During this timeframe, altered visual experience - such as dark rearing or monocular deprivation - can modulate the developmental refinement of retinogeniculate synapses ^6–9^, corticothalamic feedback ^10^, and intrinsic excitability^11^. The adult dLGN appears to be considerably less plastic, which is likely important for stable visual encoding ^9,12^. However, several examples indicate that dramatic disruptions to visual experience might trigger plasticity of dLGN and thalamocortical circuits ^13,14^, potentially similar to ways that injury or stroke in cortex opens a developmental-like period of plasticity ^15^. Such re-awakening of adult plasticity potentially permits intrinsic repair mechanisms and pharmacological approaches to activate development-like critical periods which might be a viable therapeutic approach under various disease or injury conditions ^15,16^.

Glaucoma is an ocular disease commonly associated with elevated intraocular pressure (IOP) due to reduced aqueous humor outflow from the anterior chamber ^17^. The increase in IOP damages RGC axons, ultimately leading to RGC degeneration and cutting off the route for retinal signals to reach visual centers of the brain, including the RGC-dLGN-V1 pathway ^18–20^.

We have recently mapped out the time course of this process in DBA/2J (D2) mice ^19^, a commonly-used rodent model of glaucoma that shows an age-dependent IOP elevation and RGC loss ^21–23^. These mice have a progressive IOP-associated loss of RGC axon terminals in the dLGN accompanied by functional declines in RGC-to-TC neuron synaptic transmission ^19,24^.

Thus, blindness and visual impairment arising from age-associated progressive disorders such as glaucoma might re-awaken dLGN plasticity and serve to re-shape dLGN function. Our prior work has pointed to several potentially adaptive responses in the dLGN accompanying IOP elevation and optic nerve injury. For instance, we have found enhanced action potential firing in TC neurons following experimental IOP elevation using anterior chamber microbead injections and in TC neurons from 9-month-old D2 mice ^24^. In both, measures of TC neuron excitability correlated with eye pressure, suggesting a functional link. Bilateral enucleation of adult mice likewise led to loss of RGC-TC neuron synapses accompanied by increased TC neuron intrinsic excitability ^25^. These and other unexplored effects of high IOP on TC neurons are potentially homeostatic adaptations to glaucoma in that they might allow TC neurons to continue conveying visual signals to V1 despite the diminished input from RGCs, but this possibility has not been tested.

The goal of the current study was to test whether and how glaucoma impacts TC neuron transformation of RGC synaptic inputs into action potential output. Specifically, we sought to determine whether effects on TC neurons are consistent with homeostatic compensation in response to the decline in retinogeniculate synaptic drive in aged D2 mice. We found that TC neurons from aged D2 mice generate robust synaptically-driven action potential output despite diminished retinogeniculate synaptic strength, consistent with compensatory upregulation of intrinsic excitability. This appears to be linked to augmented action potential generation mechanisms as well as diminished levels of sustained inhibition at extrasynaptic GABA_A_ receptors. These results suggest that dLGN circuits and TC neuron intrinsic function might support sustained dLGN-to-V1 transmission despite the IOP-dependent diminishment of RGC inputs to the dLGN.

## RESULTS

Intraocular pressure (IOP) is a key risk factor in glaucoma ^17^, which leads to decreased RGC signaling to the dLGN ^19^, a major thalamic relay for conscious vision. Monthly IOP measurements from DBA2J (D2) mice and their age-matched controls (D2-control) showed higher peak IOP in D2 mice indicating increased pressure on D2 retinal ganglion cells (RGCs) and their axons which make up the optic nerve (Fig. 1A). Female D2 eyes had higher peak IOP values than in males (p=0.0022, t-test), which is consistent with prior work^19,21^. To examine the optic nerve more closely, glial scarring was analyzed in cross sections of optic nerve tissue (Fig. 1B&C). Percentage of optic nerve that was glial scar was significantly higher in D2 mice indicating damage to the optic nerve and matching previous findings ^26–28^. Within the D2 population, there was more glial scarring apparent in nerves from female D2 mice compared to males, consistent with the higher IOP (p=0.018, t-test). Since there was significant glial scarring of the optic nerve in the D2 model, we tested whether RGC function was altered using pattern electroretinogram (pERG) recordings (Fig. 1D&E), finding that D2 mice showed reduced pERG P1 and N2 amplitudes compared to controls, consistent with prior findings ^29^. We did not detect an effect of sex on pERG amplitudes among D2 mice. Log-transformed P1 pERG amplitudes correlated significantly with IOP, and P1 pERG amplitudes weakly but significantly correlated with the extent of optic nerve glial scar (Fig 1. F&G). Likewise, there was a significant correlation of glial scar with IOP (R^2^=0.34, p=0.0051; Fig. S1).

**Figure 1.**
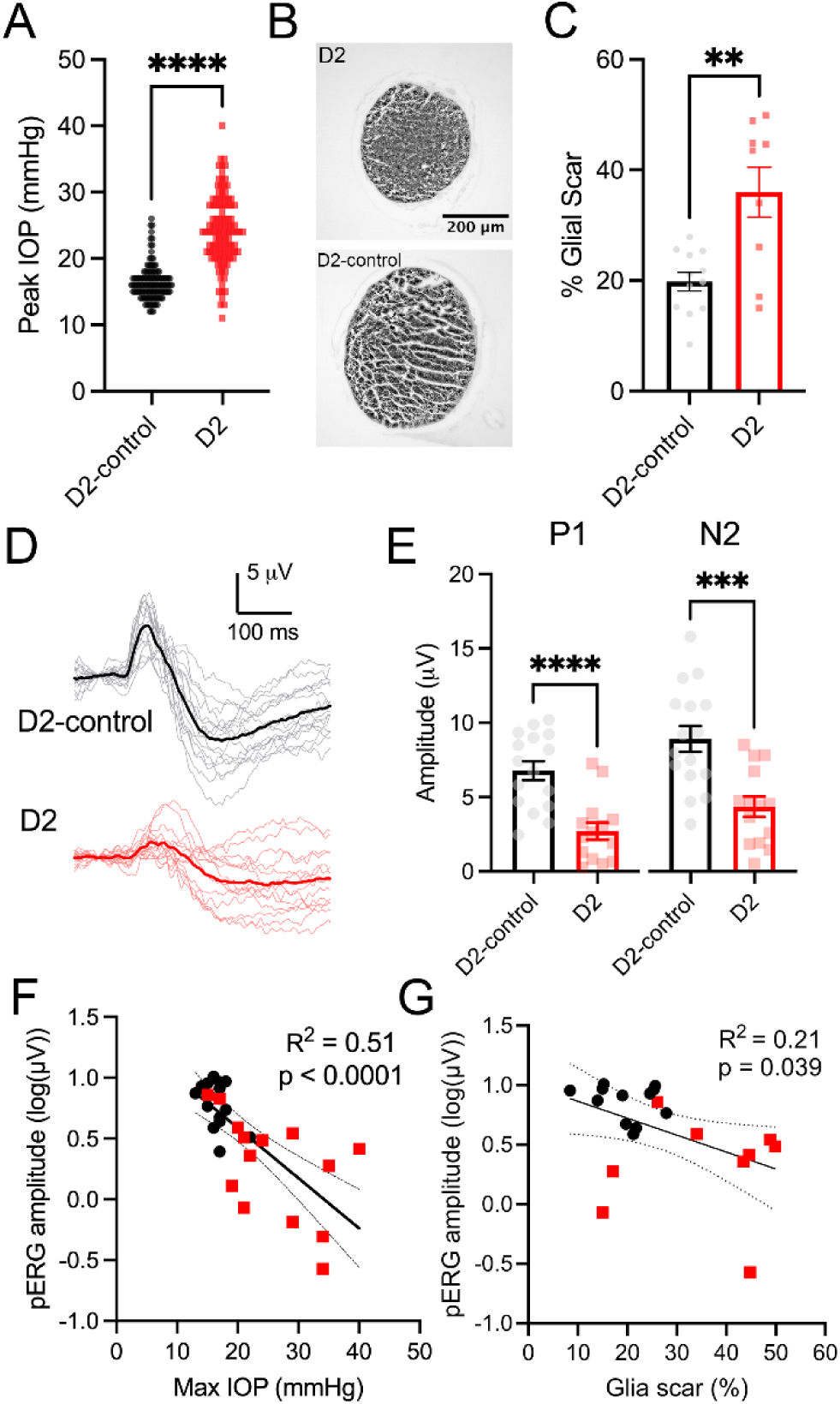
Intraocular pressure and glaucoma pathology in DBA/2J mice. A) Peak IOP measurements from individual eyes in the current study show significantly increased intraocular pressure in D2 mice compared to D2-controls [p << 0.0001 with Mann Whitney test; n = 62 mice, 124 eyes (D2-control) and n = 57 mice, 114 eyes (D2)]. B) Cross section images of optic nerves from 12-month-old D2 and D2-control animals. C) Glial scarring as a percentage of cross-sectional optic nerve area was quantified in D2 and D2-controls [p = 0.0071 with t-test; N = 12 nerves (D2-control) and N = 9 nerves (D2)]. Bars and error show mean±SEM. D) Individual (gray and light red) and average (black, dark red) pattern electroretinogram (pERG) waveforms from D2-control (black) and D2 (red) mice. E) P1 and N2 pERG amplitudes measured from pre-stimulus baseline in D2-control (black) and D2 (red) mice. P1 and N2 amplitude measurements were significantly decreased in D2 mice compared to D2-controls [(p = 0.000056 (P1) and p = 0.00033 (N2); n = 16 eyes (D2-control) and n = 13 eyes (D2)]. Bars and error show mean±SEM. F) Linear regression of P1 pERG amplitude with maximum IOP, showing a significant correlation. G) Linear regression of P1 pERG amplitude with glial scarring, showing a weak but significant correlation.

We have shown previously that D2 mice have reduced strength of RGC synaptic output to post-synaptic dLGN TC neurons ^19,24^. We therefore sought to relate how this reduction in RGC synaptic strength impacts TC neuron action potential firing (Fig 2). To accomplish this, we used a potassium-based pipette solution to switch between voltage clamp and current clamp while recording from individual dLGN TC neurons. Retinogeniculate synaptic inputs and action potential output were measured in this way in response to optic tract stimulation using a stimulus sequence derived from spiking activity of an On-sustained αRGC (Fig. 2A). Integrated charge of the excitatory postsynaptic currents (EPSCs) was reduced in D2 mice, matching the extent of reduction of EPSC amplitude (∼65%) from our prior findings ^19^. We did not detect effects of sex on EPSC amplitude within the D2 population (p=0.65, t-test). Despite the dramatic diminishment of retinogeniculate synaptic drive, the overall number of action potentials fired during the stimulus sequence was nearly identical between D2 and D2-controls across the population of recorded TC neurons (Fig. 2B). There was no significant difference between D2-control and D2 recordings for either first spike latency or first spike jitter, although the variance for both measures was higher in D2 recordings (Fig. S2). We related the action potential output to the strength of the retinogeniculate input by quantifying the total number of action potentials fired per pC of EPSC charge integrated over the stimulus sequence, finding that this ratio was higher in D2 mice compared to D2-controls (Fig. 2C). Thus, although D2 mice with increased IOP have optic nerve damage and diminished RGC drive to post-synaptic TC neurons, those TC neurons appear to more efficiently transform that decreased synaptic input into robust action potential output.

**Figure 2.**
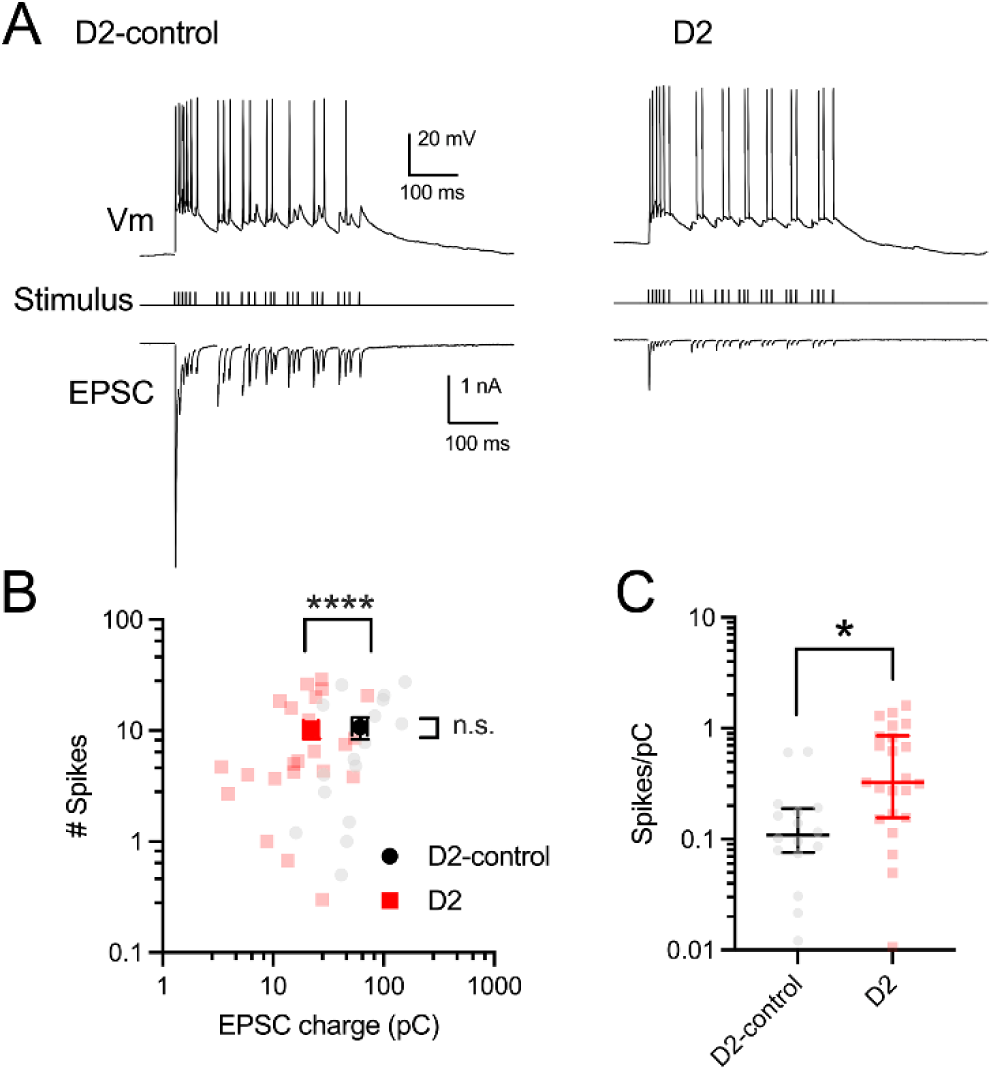
Diminished retinogeniculate synapses continue to drive robust action potential output in TC neurons from D2 mice. A) Example traces of current- and voltage-clamp recordings from TC neurons with an optic tract stimulus derived from an αRGC spike train. B) Scatterplot of EPSC charge (integrated over the stimulus train) and number of action potentials for individual cells. EPSC charge was approximately 65% lower in D2 mice compared to controls [p < 0.0001 (EPSC) via nested t-test of log transformed data] while the total number of action potentials fired over the stimulus train was similar [p = 0.92 (# of spikes) via nested t-test; n = 16 cells, 5 mice (D2-control) and n = 23 cells, 6 mice (D2)]. Group data (dark data points and error bars) are shown as mean±SEM. C) Quantification of synaptically-driven spikes per pC of EPSC charge input shows increased efficiency of synaptically-driven action potential firing in D2 mice compared to controls (p = 0.021 via nested t-test of log-transformed data). Dark lines and error bars show median±IQR.

One possible mechanism by which TC neurons in D2s might accomplish this is via enhanced intrinsic excitability. Step depolarizations and hyperpolarizations in whole cell current-clamp recordings were used to measure action potential firing and passive membrane properties of the TC neurons (Fig 3A). These experiments revealed a left-shift of the frequency-current (F-I) curves in the D2 TC neurons as well as increased incidences of depolarization block at stronger current injections (Fig. 3B&C). Passive membrane properties were measured including resting membrane potential (V_rest_), input resistance (R_in_), and membrane capacitance (C_m_; Fig. 3D-F). The mean resting membrane potential was not significantly different between D2-control and D2 mice, although there was significantly higher variance across the population of recorded D2 TC neurons (F-test p=0.0015). Within the D2 population, TC neurons from female D2 mice had a more depolarized V_rest_ than those from males (p=0.0047, t-test).

**Figure 3.**
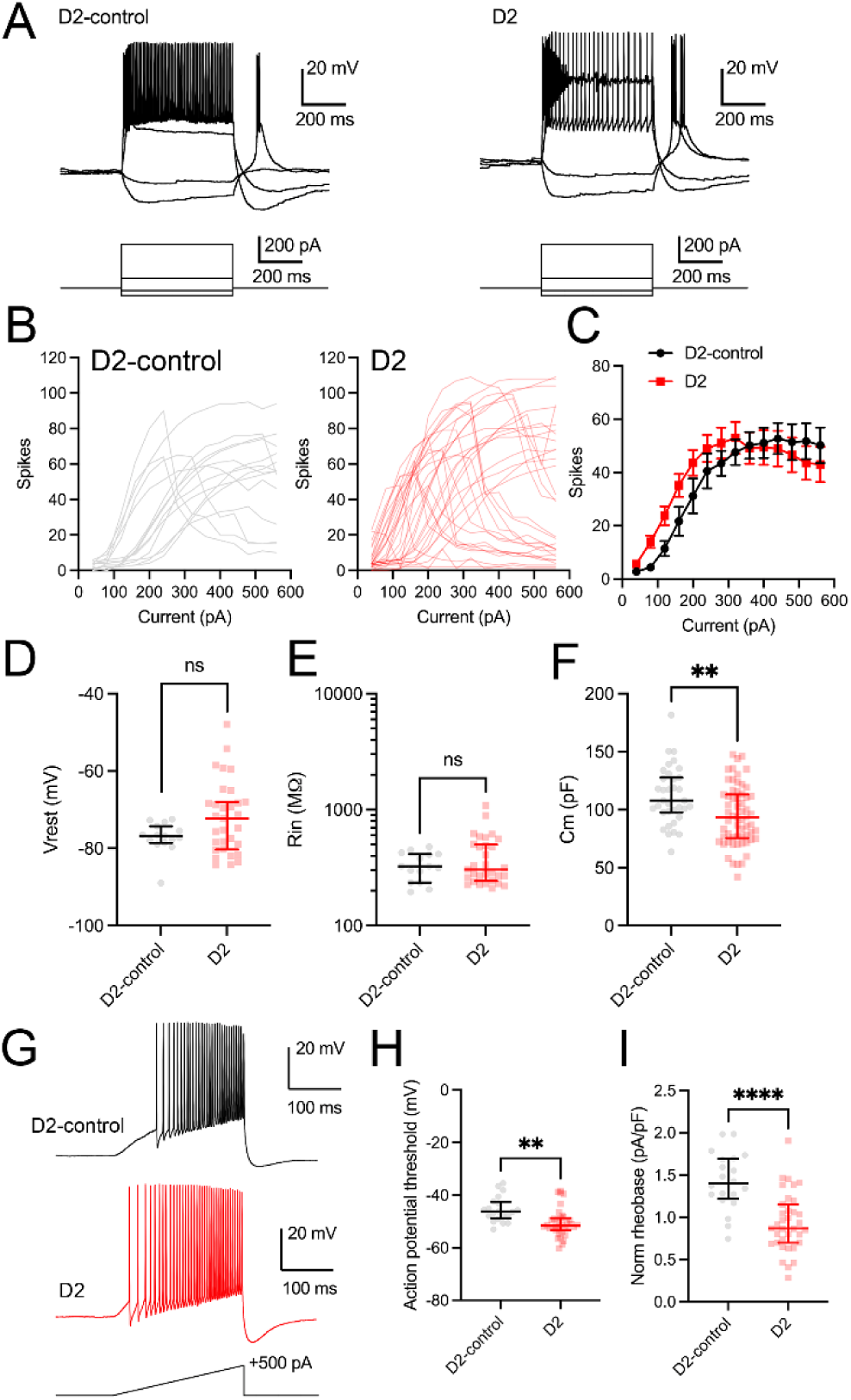
Plasticity of intrinsic excitability in D2 mice. A) Example current-clamp recordings from dLGN TC neurons in slices from D2 and D2-control mice show hyperpolarizing responses and action potential firing in response to 500-ms current injections. B) F-I plot depicting action potential numbers fired vs. injected current from individual TC neurons recorded from D2 (red) and D2-control (black) mice. C) Mean±SEM number of action potentials fired at specified currents from D2 (red) and D2-control (gray) mice. D) Resting membrane potential (V_rest_) of D2 (red) and D2-control (gray) mice shows no change between genotypes [p = 0.14 via nested t-test; n = 15 cells, 5 mice (D2-control) and n = 31 cells, 8 mice (D2)] with larger variance in D2 mice (p = 0.0015 via F-test). E) Input resistance (R_in_) measured in D2 (red) and D2-control (gray) mice shows no change between genotypes [p = 0.41 via nested t-test; n = 15 cells, 5 mice (D2-control) and n = 31 cells, 8 mice (D2)] with larger variance in D2 mice (p = 0.0021 via F-test). F) Capacitance (C_m_) measurements were lower in D2 (red) than D2-control (gray) mice [p = 0.0089 via nested t-test; n = 35 cells, 11 mice (D2-control) and n = 61 cells, 16 mice (D2)]. G) Example D2 (red) and D2-control (black) traces of current-clamp experiments using a ramp current injection (500 pA, 2 pA/ms). H) Quantification of action potential threshold shows lower first spike threshold in D2 (red) than in D2-control (black) mice in response to the ramp depolarization [p = 0.014 via nested t-test; n = 25 cells, 10 mice (D2) and n = 20 cells, 6 mice (D2-control)]. I) Rheobase, measured during ramp depolarization, was lower in D2 (red) compared to D2-control (black) mice [p = 0.00018 via nested t-test; n = 35 cells, 10 mice (D2) and n = 20 cells, 6 mice (D2-control)]. Dark lines and error bars in panels D-F, H, and I show median±IQR.

Hyperpolarizing steps were used to measure the input resistance, and there was no significant difference between D2-control and D2 mice, although variance was significantly higher in the D2 group (F-test, p=0.0021). R_in_ was higher in TC neurons from D2 females compared to those from D2 males (p=0.012, t-test). C_m_ was measured in voltage clamp recordings by integrating the membrane capacitance transient in response to a −10 mV step. C_m_ was significantly lower in D2 TC neurons, consistent with a decreased TC neuron membrane surface area. There was a difference in C_m_ between TC neurons from male vs female D2 mice, with those from females having a lower C_m_ (p=0.015, t-test). We next studied action potential properties while maintaining resting potential at approximately −60 mV to inactivate low voltage-activated Ca^2+^ currents ^30^, which contribute to TC neuron rebound spiking ^31–33^. Using a 500 pA ramp current injection (Fig. 3G-I), we measured TC neuron action potential threshold and rheobase, finding that TC neurons from D2 mice had a lower action potential threshold and lower rheobase. Threshold was slightly lower in TC neurons from D2 females compared to males (p=0.029, t-test), although we did not detect a sex difference in rheobase (p=0.31).

Overall, these data indicate that TC neurons from D2 mice are more excitable as indicated by 1) an increased readiness to fire action potentials in response to stimulation, 2) the increased incidence of depolarization block, 3) the shift in action potential threshold, and 4) a reduction in rheobase. Using current clamp to maintain resting potential indicates that the increased excitability is at least partially independent of effects on V_rest_.

The axon initial segment (AIS) is the structure responsible for initiating action potentials due to its high density of voltage-gated sodium channels ^34,35^. AIS length can be modulated in some neuron populations as an apparent mechanism to homeostatically regulate intrinsic excitability ^36,37^. IOP-induced alterations in AIS length in mice with glaucoma might underlie changes in TC neuron excitability in D2 mice. To test this, we performed immunostaining for Ankyrin-G, a scaffolding protein found in the AIS, to determine the size of the AIS in TC neurons (Fig. 4A&B). We manually measured the AIS length in images obtained from Ankyrin-G-stained dLGN slices. Overall, AIS length was similar and not significantly different when comparing between measurements from D2 and D2 control dLGN sections. Thus, modulation of AIS length in dLGN TC neurons is unlikely to underlie their altered excitability in D2 mice.

**Figure 4.**
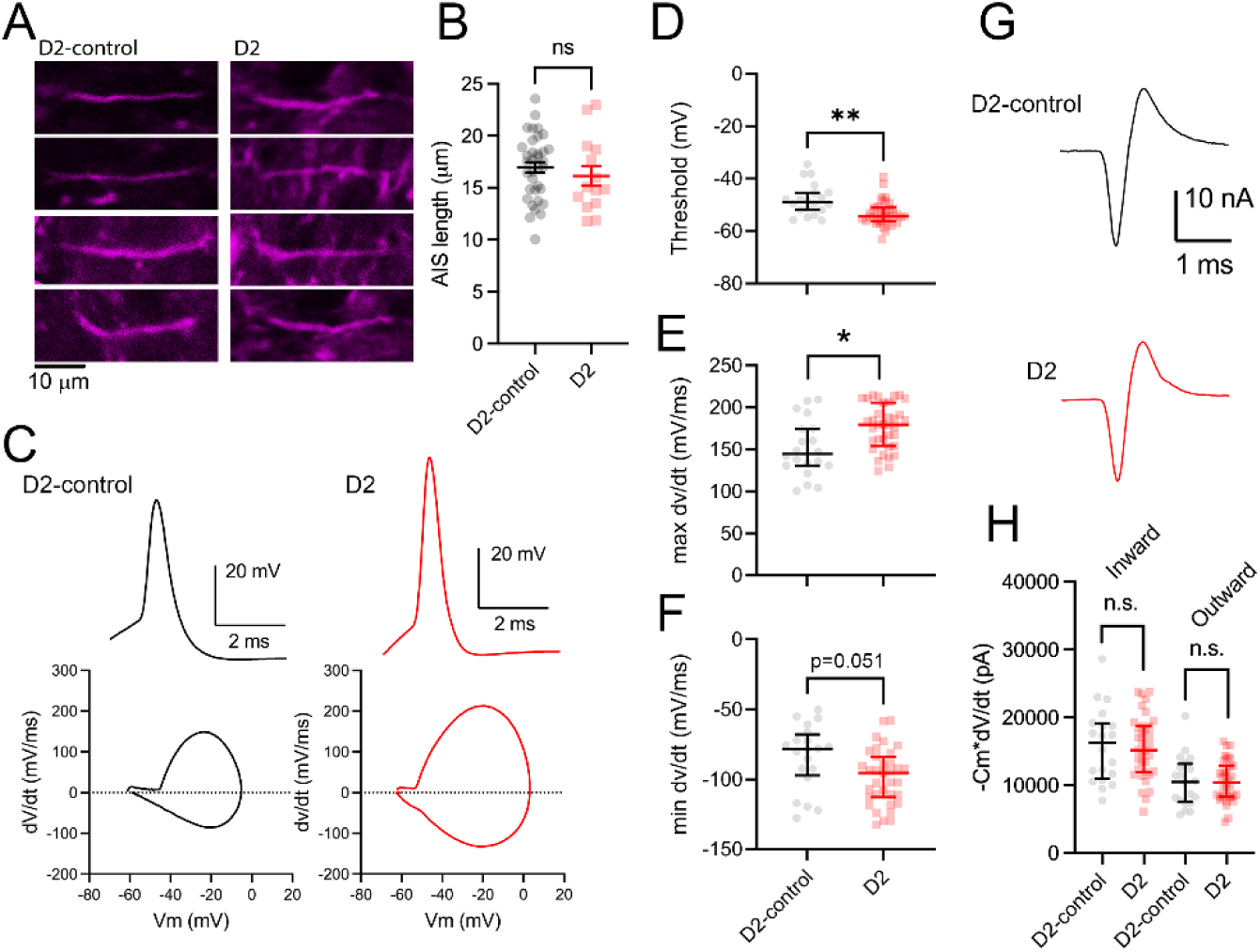
D2 action potential waveforms are altered indicating increased Na^+^ and K^+^ current density. A) Example images of D2 and D2-control dLGN stained for ankyrin-G to identify axon initial segments. B) Average axon initial segment length for individual mice shows no change between D2 (red) and D2-control (black) [p = 0.32 with nested t-test; n = 1715 AISs, 35 mice (D2-control) and n = 707 AISs, 14 mice (D2)]. Dark lines and error bars show mean±SEM. C) Example action potential waveforms and action potential phase plots (dv/dt) recorded from TC neurons from D2 (red) and D2-control (black) mice. D) Analysis of action potential threshold from phase plots shows lower threshold in D2 mice [p = 0.0064 with nested t-test; n = 20 cells, 6 mice (D2-control) and n = 35 cells, 10 mice (D2). E) Analysis of maximum dv/dt shows an increase in D2 mice [p = 0.018 with nested t-test; n = 20 cells, 6 mice (D2-control) and n = 35 cells, 10 mice (D2)]. F) Analysis of minimum dv/dt shows no significant difference between genotypes [p = 0.051 with nested t-test; n = 20 cells, 6 mice (D2-control) and n = 35 cells, 10 mice (D2)]. Dark lines and error bars in panels D-F show median±IQR. G) Example traces of membrane currents underlying the action potential (calculated as - Cm*dv/dt) in D2 (red) and D2-control (black) dLGN. H) Analysis of -Cm*dv/dt shows no significant difference in the amount of current charging the membrane [p = 0.87 (inward) and p = 0.97 (outward) with nested t-test] between D2 and D2-control TC neurons. Dark lines and error show median±IQR.

To further examine the relationship between the TC neurons and action potential generation, we analyzed action potential waveforms via phase plots ^38^. Taking the first derivative of the action potential waveform and plotting it against voltage allows us to examine different features of the action potential (Fig. 4C-F). We found a hyperpolarized shift in action potential threshold in TC neurons from D2 mice, consistent with experiments using the ramp stimulus, above. There was also a significant increase in peak slope of the rising phase of action potentials in D2 TC neurons (max dv/dt) with a trending but not statistically significant effect (p=0.051) on the peak slope of the falling phase (min dv/dt). This indicates that TC neurons in D2 mice seem to show changes in excitability that correlate with changes in the density of voltage gated sodium channels. Next, we looked at current flow responsible for charging the membrane (-C_m_*dv/dt) during TC neuron action potentials (Fig. 4G&H) ^39,40^. Both the inward and outward current amplitudes were comparable between D2-control and D2 mice, with no significant difference between groups. We did not detect a significant difference by sex for any of these parameters within the D2 population. Together, these results indicate that although the total membrane current is the same, the lower C_m_ (Fig. 3F) of D2 TC neurons leads to an increased current density of Na^+^ currents, likely contributing to enhanced action potential generation.

In addition to effects on intrinsic excitability, altered synaptic properties such as reduced synaptic inhibition might support TC neuron action potential firing in response to excitatory input. To test this possibility, TC neuron excitatory and inhibitory post-synaptic currents (EPSCs and IPSCs) were measured in response to a physiological train of optic tract stimulation similar to that used in Fig. 2, except using a Cs-based pipette solution (Fig. 5A). In this approach, EPSCs represent monosynaptic excitatory drive from RGC axons while the IPSCs are the result of disynaptic feed-forward inhibition arising from local interneurons ^1,41^. We found that EPSCs in brain slices from D2 mice were reduced in amplitude compared to D2-controls, as above (Fig. 2) and in keeping with our prior findings ^19^. IPSCs from D2-controls were relatively sustained but showed a notably faster decay back to baseline in slices from D2 mice. To quantify this, we used a single stimulation of the optic tract while recording IPSCs (Fig. 5B). These IPSCs were entirely blocked by 25 μM SR95531 (n=6 cells, 100±2% reduction in amplitude, Fig. S3) indicating appropriate voltage clamp isolation of inhibitory currents. The decay of the IPSCs could be fit with a sum of two exponential functions, and while the time constants themselves were not significantly different between D2 and D2-control TC neurons, there was a relative decrease in the percentage of the decay mediated by the slow time constant (Fig. 5C&D) and this did not differ by sex within the D2 population (p=0.6, t-test).

**Figure 5.**
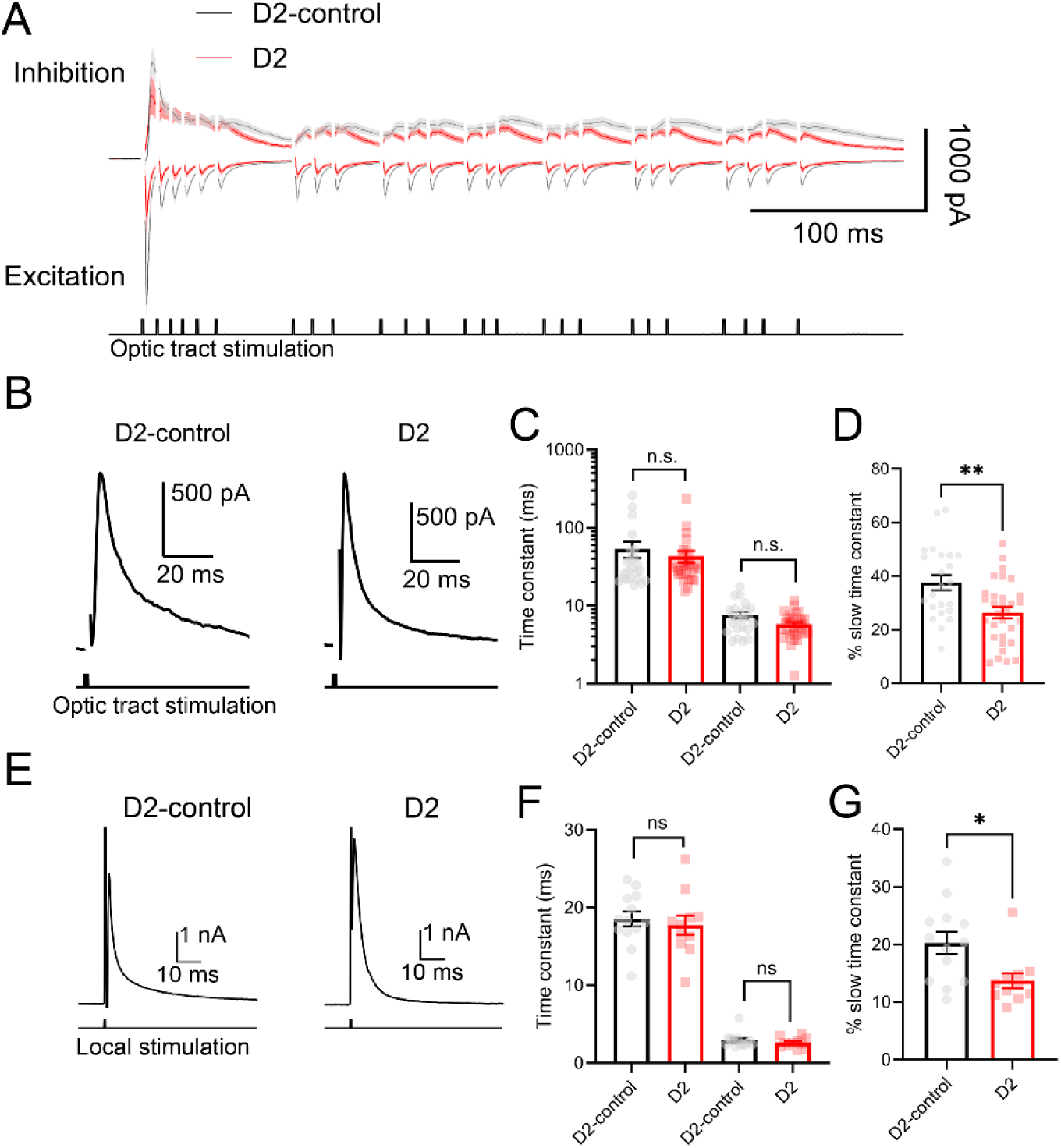
Feedforward inhibition is more transient in the D2 mouse. A) Mean±SEM traces of monosynaptic retinogeniculate excitatory post-synaptic currents (EPSCs, “Excitation”) and feed-forward disynaptic inhibitory post-synaptic currents (IPSCs, “Inhibition”) driven by optic tract stimulation and recorded from TC neurons in D2 (red) and D2-control (gray) dLGN slices [n = 9 cells from 3 mice (D2-control) and n = 23 cells from 7 mice (D2)]. B) Example feed-forward IPSC traces following single optic tract stimulation illustrating faster decay kinetics in the D2 recording. C) Decay time constants show no difference between D2 (red) and D2-control (gray) mice following single optic tract stimulation [p = 0.063 (fast time constant) and p = 0.65 (slow time constant) via nested t-test of log transformed values; n = 31 cells, 9 mice (D2) and n = 23 cells, 9 mice (D2-control)]. D) Percentage of decay made up of slow time constant is lower in D2 mice (red) compared to controls (gray) following optic tract stimulation. [p = 0.0020 via nested t-test; n = 31 cells, 9 mice (D2) and n = 23 cells, 9 mice (D2-control)]. E) Example IPSC traces in response to stimulation delivered by an aCSF-filled pipette located approximately 25 microns from the recorded TC neuron in the presence of CNQX and D-AP5. F) Decay time constants show no difference between D2 (red) and D2-control (gray) mice following local stimulation. [p = 0.41 (fast time constant) and p = 0.62 (slow time constant) via nested t-test; n = 11 cells, 7 mice (D2) and n = 13 cells, 8 mice (D2-control)]. G) Percentage of decay made up of the slow time constant is lower in D2 mice (red) compared to controls (gray) following local stimulation [p = 0.029 via paired t-test; n = 11 cells, 7 mice (D2) and n = 13 cells, 8 mice (D2-control)]. Bar graphs and error bars show mean±SEM.

Although this stimulus approach allowed us to record inputs from local interneurons, it is potentially confounded by glaucomatous alterations of RGC-to-interneuron synaptic function. Therefore, we recorded monosynaptic IPSCs while bypassing the RGCs by directly stimulating interneurons with an extracellular electrode positioned in the dLGN approx. 25 microns from the recorded TC neurons (Fig. 5E). The aCSF was supplemented with glutamatergic blockers (CNQX, D-AP5) to further isolate inhibitory inputs in the absence of RGC inputs. IPSCs were completely blocked by the GABA_A_ receptor blocker SR95531 (25 μM, n=10 cells; 99±1% reduction) and unaffected by application of the GABA_B_ receptor blocker CGP55845 (0.81+/-5% reduction, n = 6 cells; Fig. S4). Similar to above, the decay of IPSCs recorded in this way was well-fit with a sum of two exponential functions. Although fast and slow time constants were similar between D2 and D2-control recordings, the D2 recordings showed a reduction in the relative percentage of the decay mediated by the slow time constant (Fig. 5F&G) and this was more pronounced in TC neurons from female D2 mice compared to male D2 mice (p=0.032, t-test). Overall, these results show that feedforward inhibition is more transient in TC neurons from D2 mice. This effect on sustained inhibition likely contributes, alongside the enhanced intrinsic excitability, to enhanced TC neuron action potential generation during optic tract stimulation.

We next employed pharmacological manipulations to investigate GABAergic signaling in the dLGN. Within the thalamus, GABA acts extrasynaptically on δ subunit-containing GABA_A_ receptors ^42–44^. The relative reduction in the slow component of the IPSC decay might be the result of diminished extrasynaptic GABA effects ^45,46^. We therefore tested whether DS2, a positive allosteric modulator of δ subunit-containing GABA_A_ receptors ^47^, had differential effects on TC neuron IPSCs from D2 vs D2-control mice (Fig 6A). DS2 (10 μM) evoked an outward current of 171±25 pA in D2-control TC neurons and 119±7 pA in D2 TC neurons, but this difference was not statistically significant (p=0.061, nested t-test; Fig. S5). We again recorded IPSCs from dLGN TC neurons in response to local extracellular stimulation in the presence of CNQX and D-AP5 and found that DS2 enhanced the slower components of the IPSC. This effect was more pronounced in recordings from D2-controls compared to those from D2 mice (Fig. 6B), implying a reduced contribution of extrasynaptic GABA_A_ receptors in D2 TC neurons.

**Figure 6.**
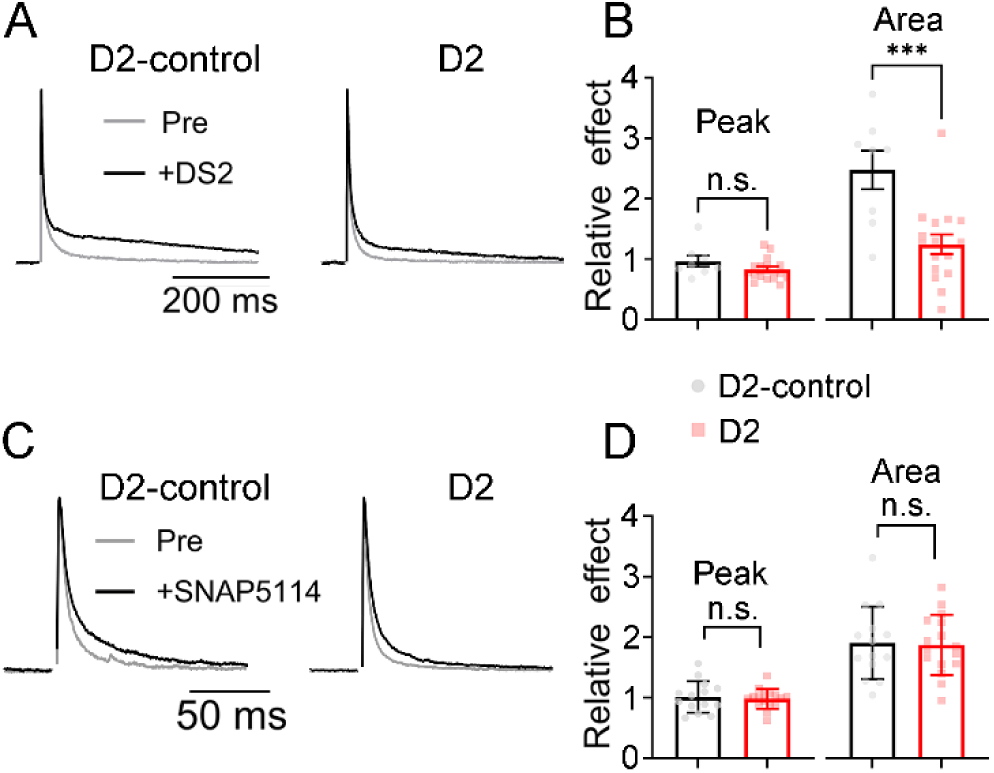
Pharmacology of inhibition indicates decreased activation of extrasynaptic GABA_A_ receptors in DBA/2J TC neurons. A) Example peak-normalized IPSCs from TC neurons from D2 and D2-control mice evoked by stimulation with an aCSF-filled patch pipette positioned approximately 25 microns from the recorded TC neuron in the presence of CNQX (20 μM) and D-AP5 (50 μM). Traces show before (gray) and after (black) bath application of 10 μM DS2. B) Relative effect of DS2 application on peak response of the IPSC (p = 0.16 via nested t-test) and area (p = 0.0010 via nested t-test) of D2 (red) and D2-control (black) mice [n = 8 cells, 4 mice (D2-control) and n = 16 cells, 6 mice (D2)]. C) Example peak-normalized traces of IPSCs from D2 and D2-control mice before (gray) and after (black) application of SNAP5114 (60 μM). IPSCs were evoked as in A. D) Relative effect of SNAP5114 application on peak response (p = 0.70 via nested t-test) and area (p = 0.91 via nested t-test) of D2 (red) and D2-control (black) mice [n = 16 cells, 6 mice (D2) and n = 14 cells, 5 mice (D2-control)]. Bar graphs and error bars show mean±SEM.

We next investigated whether this could be attributable to differential GABA reuptake by astrocytes in D2 vs. D2-control dLGN. To test this, we applied the GABA transporter inhibitor SNAP5114 (60 μM), which is fairly selective for the astrocytic GAT-3 transporter ^48–50^, while again measuring IPSCs evoked by local stimulation (Fig. 6C). SNAP5114 had no effect on IPSC peak amplitude but led to an increase in IPSC area due to enhancement of later portions of the IPSC. However, the relative change in the sustained portion of the IPSC by SNAP5114 was similar in D2 and D2-control recordings (Fig. 6D), implying that astrocytic GABA uptake is similar in control mice and mice with glaucoma. Together, these results suggest a decrease in extrasynaptic inhibition in dLGN TC neurons of D2 mice, likely supporting enhanced action potential firing in response to diminished excitatory synaptic drive from RGCs.

## DISCUSSION

In healthy mouse dLGN, 1-3 RGC inputs, largely arising from the same class of RGC, provide the major excitatory input driving TC neuron spike output, with a threshold of approximately 600 pA EPSC needed to trigger a TC neuron action potential ^51–53^. A loss of retinogeniculate synaptic input in the glaucomatous dLGN ^19^ would therefore be predicted to diminish the efficiency of TC neuron action potential output. However, the major finding of this study is that dLGN TC neurons in 11-15 month-old D2 mice remain able to efficiently transform retinal ganglion cell excitatory synaptic inputs into action potential output despite the diminished strength of those inputs. This occurs at an age where these mice show clear signs of glaucoma including high IOP, optic nerve pathology, and declining pERG responses, among numerous other pathological signs documented in previous studies by our group and others ^19,21,24,26–29,54–61^. We attribute this to a combination of increased intrinsic excitability - likely the result of an interplay between both increased voltage-gated channel density and effects on passive membrane properties – and a reduced extrasynaptic inhibition from GABA spillover. In several measured parameters, we detected differences between recordings from TC neurons from male vs. female D2 mice, which might be partially attributable to the higher overall IOP in female D2 mice^19,21^. Future studies should be designed and powered to specifically examine sex differences within the D2 population. The fact that TC neurons can maintain the same levels of action potential output despite an approximately 65% decline in RGC input strength suggests that these might be compensatory/homeostatic responses to glaucoma pathology.

The current study also sought to distinguish the features of the enhanced TC neuron intrinsic excitability in D2 mice - whether it is simply the result of passive properties such as increased input resistance and depolarized resting potential or could be attributed to active spike generation mechanisms. Notably, while we did not find a significant difference in V_rest_ or R_in_ between D2 and D2-controls, similar to what we showed previously in younger (9-month-old) D2 mice ^24^, both measures were significantly more variable in D2 recordings, suggesting considerable heterogeneity across the population of TC neurons, even within individual mice. The effects on resting membrane potential were clearly not the sole contributor to TC neuron action potential generation, as experiments where we controlled V_rest_ using stable current injections revealed lower action potential threshold and rheobase in D2 mice.

The lower threshold and rheobase might suggest plasticity of TC neuron axon initial segments, which are densely packed with Na^+^ channels and critical for action potential generation. AIS length can be a locus for plasticity of intrinsic excitability in neurons in other brain regions ^35,37,62,63^. In dLGN TC neurons, the concentration of K^+^ channels in the AIS has been shown to be associated with developmental plasticity of TC neuron excitability ^11^. Here, AIS length was similar between D2 and D2-control mice. Analysis of action potential waveforms and the membrane currents responsible for charging the TC neuron membrane during the action potential showed that while the absolute amplitudes of currents were the same when comparing D2 and D2-control recordings, the D2 TC neurons had a smaller membrane capacitance, leading to a higher current density.

The contributions of intrinsic action potential generation mechanisms to support synaptically-driven spiking by TC neurons appeared to be complemented by alterations in extrasynaptic inhibition. In addition to local synaptic inhibition, dLGN TC neurons show considerable tonic inhibition and apparent GABA spillover from local interneurons that is attributable to activation of high-affinity δ subunit-containing extrasynaptic GABA_A_ receptors ^42–44,47,64,65^. While there is no spillover component of single-vesicle quantal IPSCs ^44^, our results show that DS2 enhanced a slow component of the multiquantal IPSC, consistent with GABAergic spillover. During a train of RGC inputs, feedforward inhibition from local interneurons generates a sustained inhibitory response in recorded TC neurons that is notably diminished in TC neurons from D2 mice. Different IPSC kinetics - with D2 TC neurons showing faster IPSC decays - along with differential effects of DS2 on the sustained component of the IPSC indicate a reduced activation of extrasynaptic GABA receptors in D2 TC neurons, which would allow the diminished excitatory inputs to have bigger effects on TC neuron voltage. This is likely to be the result of fewer extrasynaptic receptors rather than increased GABA uptake by astrocytes as SNAP5114 ^48^, a GABA transporter inhibitor, had led to a similar extent of enhancement of the sustained component of IPSCs in D2 and D2-controls.

Enigmatically, we have found previously that D2 mice show opposite effects on synaptic inhibition ^66^ compared to the effects on extrasynaptic inhibition here. In ∼12 month-old D2 mice, TC neurons have an apparent increase in synaptic GABA receptors, which leads to overall larger quantal IPSC amplitudes and larger local interneuron-driven multiquantal IPSCs than would be expected based on the extent of RG synapse loss. This seems at odds with a traditional “homeostatic” response to diminished excitatory drive from RGCs whereas the reduction in extrasynaptic inhibition seems to more clearly align with a mechanism for boosting action potential firing. Some evidence in other systems indicates that synaptic and extrasynaptic GABA receptor populations exist in dynamic competition with one another ^67,68^. Still, we do not know what effect the combined increase in synaptic inhibition and decrease in extrasynaptic inhibition ultimately has on TC neuron output or “why” a TC neuron might boost synaptic inhibition while simultaneously diminishing the extrasynaptic response to GABA spillover. One possibility is that changing the balance of both forms of inhibitory input could contribute to TC neuron input selectivity or spiking precision in the face of weakened retinogeniculate inputs ^41,43,69–71^. In the dLGN, feed-forward inhibition that is “locked” to the excitatory input, meaning that it arises from local interneurons driven by the same RGC axons driving the recorded TC neuron at the retinogeniculate glomerulus, increases spike precision and restricts the TC neuron to fire a single post-synaptic spike per presynaptic spike ^41^. This is especially the case in TC neurons operating in a tonic firing mode (rather than burst mode, at more negative V_rest_). In our recordings of synaptically-driven spiking, first spike latency and first spike jitter were slightly higher in TC neurons from D2 mice although the difference compared to D2-controls was not significant. This might be indicative of intrinsic and/or synaptic mechanisms preserving spike precision, but whether this is actually the case will need further study - possibly with pharmacological or modeling approaches. In contrast, inhibition arising from RGC activation by RGCs not inputting onto a recorded TC neuron has been suggested to play a role in shaping TC neuron receptive field properties ^72–74^. This also could be modulated by the shifted balance of phasic/sustained inhibition and merits future study.

Our results have implications for the broader understanding of plasticity in the adult dLGN. Overall, the picture that emerges is one of homeostasis, with multiple cell-intrinsic and synaptic properties in dLGN TC neurons adapting to support synaptically-driven action potential generation even as those synaptic inputs from RGCs decline due to RGC degeneration.

Notably, we have not found evidence for homeostatic synaptic scaling on excitatory synapses with declining retinogeniculate synaptic strength, as mEPSCs and AMPA/NMDA ratios were comparable between DBA/2J and controls in our prior studies ^19,24^. This is in contrast to developmental plasticity, which involves AMPA trafficking at retinogeniculate synapses ^52,75^.

Additionally, early monocular deprivation can impact corticothalamic feedback synapses, apparently via both pre- and post-synaptic mechanisms ^10^. Instead, in the aged mouse dLGN, glaucoma pathology appears to trigger effects on both inhibitory synapses and extrasynaptic sites as well as on TC neuron intrinsic excitability. There are relatively few examples of plasticity in the adult dLGN, with most research being focused on experience-dependent plasticity during development. One study using longitudinal in vivo imaging of TC neuron axon terminals in binocular V1 following a brief monocular deprivation in adult mice showed an increase in response strength to stimulation of the non-deprived eye ^76^. This was a novel finding in that it identified a thalamic locus for some adult ocular dominance plasticity, which was thought to reside solely in visual cortex with minimal capacity for plasticity in the mature dLGN. Our findings of apparently homeostatic compensation in the adult glaucomatous dLGN add support to the notion of adaptive plasticity in the adult dLGN. This might reflect an awakening of critical period-like plasticity triggered under disease conditions, although the effects clearly differ from those seen during dLGN maturation over the first few postnatal weeks in mice.

While it is likely that high IOP and loss of RGC synapses are the ultimate triggers for the effects on dLGN inhibition and TC neuron excitability, we do not know what transduces these effects. Classically, Ca^2+^-dependent transduction pathways involving calcium/calmodulin-dependent kinases serve as the “thermostat” to detect altered activity levels and trigger intrinsic excitability changes and/or synaptic scaling – either throughout a neuron or at specific synapses^77^. Glial cells, which are constantly surveilling the brain environment, might also contribute by detecting altered activity levels and releasing diffusible signals such as neurotrophins (i.e. BDNF) or cytokines (i.e. TNFα) that can trigger synaptic or intrinsic plasticity via transduction pathways activated by their respective receptors such as the Trk family and p75NTR receptors for neurotrophins and a wide array of cytokine receptors ^78–87^. We have recently shown that dLGN microglia respond to elevated IOP in DBA/2J mice and might take on a protective phenotype ^88^, possibly supporting TC neuron compensatory responses. Overall, the cell-intrinsic or local mechanisms causing high IOP/glaucoma to trigger TC neuron compensatory responses are unknown and merit future research.

## METHODS

| REAGENT or RESOURCE | SOURCE | IDENTIFIER |
| --- | --- | --- |
| **Antibodies** |  |  |
| Anti-ankyrin G, rabbit polyclonal | Synaptic Systems | Cat # 386 003, RRID:AB_2661876 |
| Anti-rabbit, donkey IgG, Alexa Fluor 568 | Invitrogen | Cat # A-10042, RRID:AB_2534017 |
| **Experimental organisms** |  |  |
| DBA/2J mouse ("D2") | The Jackson Laboratory | Strain: 000671, RRID:IMSR_JAX:000671 |
| DBA/2J-Gpnmb<sup>+</sup>/SjJ ("D2-control") | The Jackson Laboratory | Strain: 007048, RRID:IMSR_JAX:007048 |
| **Software** |  |  |
| ImageJ/FIJI | Fiji, https://fiji.sc | RRID:SCR_002285 |
| Prism 10 | GraphPad | RRID:SCR_002798 |
| pClamp | Molecular Devices | RRID:SCR_011323 |
| **Chemicals** |  |  |
| CNQX | Tocris | Cat # 0190 |
| D-AP5 | Tocris | Cat # 0106 |
| SR95531 | Tocris | Cat # 1262 |
| DS2 | Tocris | Cat # 3679 |
| (S)-SNAP5114 | Tocris | Cat # 1561 |

### Mice and IOP Measurements

All animal procedures were reviewed and approved by the Institutional Animal Care and Use Committee (IACUC) at the University of Nebraska Medical Center. DBA/2J mice (D2, The Jackson Laboratory #000671, RRID:IMSR_JAX:000671) and DBA/2J-*Gpnmb1^+^*mice (D2-control, The Jackson Laboratory #007048, RRID:IMSR_JAX:007048) were used for this study at 11-15 months of age. Mice of both sexes were used. The mice were bred and housed at the University of Nebraska Medical Center Comparative Medicine facilities on a 12h/12h light-dark cycle with free access to food and water. Intraocular eye pressure measurements (IOPs) were taken on D2 and D2-control eyes initially beginning at 3 - 5 months of age and subsequent IOPs were measured approximately monthly throughout the study. IOPs were obtained using a TONOLAB rebound tonometer (iCare, Finland), displayed in mmHg. D2 and D2-control mice were lightly anesthetized with inhaled isoflurane (Piramal Critical Care, Bethlehem, PA). Mice were put in an induction chamber with approximately 4% isoflurane until sedated. Approximately 2% isoflurane, administered through a nose cone, was used to maintain sedation while six consecutive, low-error IOP measurements were taken from the central cornea to generate a single readout. Three readouts per eye were averaged to provide a single IOP value per eye for each time-point.

### Pattern Electroretinogram (pERG)

Mice were dark-adapted overnight prior to pERG recordings, which were performed using the Celeris Small Animal ERG system (Diagnosys LLC, Lowell MA USA). An intraperitoneal ketamine/xylazine injection [100 mg/kg ketamine (Zoetis, Parsipanny, NJ) and 5 mg/kg xylazine (Akorn Inc., Lake Forest, IL)] was used to anesthetize the mice, and tropicamide mydriatic eye drops (1%, Alcon Laboratories, Fort Worth, TX) and proparacaine topical anesthetic eye drops (0.5% Alcon Laboratories, Fort Worth, TX) were applied to each eye. Once the mouse was anesthetized, hypromellose gel (0.3%; GenTeal Tears, Alcon, Fort Worth, TX) was applied to the eyes, and an electrode/fiber optic stimulator was placed on one eye to serve as a reference electrode while a pattern stimulator/electrode was positioned on the other. The pattern stimulator was then used to deliver a 100% contrast reversing horizontal bar stimulus (0.059 cycles/degree, 50 cd/m^2^, 2.1 Hz pattern reversal) while pERG waveforms were acquired at 2 kHz and bandpass filtered at 1-50 Hz. After acquiring pERG data from one eye, electrodes were switched, and pERG data were recorded from the other eye.

### Slice Preparation and Electrophysiology

For preparation of slices for electrophysiology, the “protected recovery” method was used ^89,90^. After mice were killed via CO_2_ inhalation and cervical dislocation, brains were extracted and submerged in a slush of artificial cerebral spinal fluid (aCSF) for ∼ 1 min. The aCSF was comprised of the following: 128 mM NaCl, 2.5 mM KCl, 1.25 mM NaH_2_PO_4_, 24 mM NaHCO_3_, 12.5 mM glucose, 2 mM CaCl_2_, and 2 mM MgSO_4_. For coronal slices containing dLGN, cerebellum was removed and the flat, exposed surface glued to the platform of a Leica VT1000S vibratome for slicing (250 micron thickness). For parasagittal slices containing optic tract and dLGN, the brain was hemisected ∼5 degrees from the medial longitudinal fissure and ∼20 degrees from the horizontal plane ^91,92^, after which the medial surface was glued to the platform for slicing (300 micron thickness). After slicing, tissue was incubated for 12 minutes in a warmed (32°C) N-methyl-D-glucamine (NMDG) solution (92 mM NMDG, 2.5 mM KCl, 1.25 mM NaH_2_PO_4_, 25 mM glucose, 30 mM NaHCO_3_, 20 mM HEPES, 0.5 mM CaCl_2_, 10 mM MgSO_4_, 2 mM thiourea, 5 mM L-ascorbic acid, and 3 mM Na-pyruvate and bubbled with 95% O_2_ and 5% CO_2_). Both the aCSF and NMDG solutions were prepared with a pH of 7.4 and osmolality of 300 – 315 mOsm. Slices were allowed to recover for at least 1 hour in room temperature aCSF (bubbled with 95% O_2_/5% CO_2_) prior to recording. Patch pipettes were pulled from thin-walled borosilicate glass with an internal filament and filled with either a Cs^+^-or K^+^-based pipette solution (Cs^+^-based solution: 120 mM Cs-methanesulfonate, 2 mM EGTA, 10 mM HEPES, 8 mM TEA-Cl, 5 mM ATP-Mg, 0.5 mM GTP-Na_2_, 5 mM phosphocreatine-Na_2_, 2 mM QX-314; K+-based solution: 120 mM K-gluconate, 8 mM KCl, 2 mM EGTA, 10 mM HEPES, 5 mM ATP-Mg, 0.5 mM GTP-Na_2_, 5 mM phosphocreatine, *pH* = 7.4, 275 mOsm). Reported voltages are corrected for liquid junction potentials (10 mV for Cs^+^ and 14 mV for K^+^ pipette solutions). Electrophysiology data were acquired at 10 kHz with an Axon MultiClamp 700B amplifier and Digidata 1550 Digitizer (Molecular Devices). While recording, slices were superfused with aCSF (2 mL/minute) that was bubbled with 5% CO_2_ / 95% O_2_ and warmed to 30-33°C with an in-line solution heater. Excitatory and inhibitory post-synaptic currents (EPSCs and IPSCs) evoked by optic tract stimulation were recorded while holding the TC neurons at −70 mV or −74 mV (for recording EPSCs with Cs^+^ or K^+^ pipette solutions, respectively) and 0 mV (for recording IPSCs), in response to stimulation of the optic tract using a aCSF-filled patch pipette (100-400 μA, 0.2 ms stimulus duration using an A-M Systems Isolated Pulse Stimulator) while synaptically-driven spiking was recorded in current clamp configuration. For recording locally-driven IPSCs ^93^, an aCSF-filled patch pipette was positioned in the extracellular space approximately 25 microns from the recorded TC neuron in coronal slices and used to deliver a current stimulus (0.2-0.3 ms, 100-400 μA) in the presence of glutamatergic blockers (20 μM CNQX, 50 μM D-AP5).

Intrinsic excitability was monitored in current clamp. A series of depolarizing current injections (500 ms duration) was used to measure the relationship of action potential firing to depolarizing current injection. For measuring rheobase, V_m_ was maintained at approximately –60 mV to inactivate LVA Ca^2+^ currents ^30^ and a ramp current stimulus (500 pA amplitude, 2 pA/ms) was delivered. Rheobase was taken as the current stimulus at the threshold of the first action potential. To generate action potential phase plots and estimate membrane currents responsible for action potential generation, TC neurons were stably depolarized to –60 mV using a DC injection, and a short (2 ms) depolarizing current injection was used to evoke an action potential. Membrane capacitance was measured by integrating the whole-cell capacitance transient current over 10 ms in response to a –10 mV hyperpolarizing voltage step. Input resistance was measured in current clamp in response to a series of hyperpolarizing current injections. For constructing phase plots, the sampling frequency was increased to 100 kHz. For pharmacologically manipulating dLGN inhibition, we bath-applied DS2 (10 μM), a positive allosteric modulator of δ-subunit-containing GABA_A_ receptors, or the GABA transporter inhibitor SNAP5114 (60 μM), which disrupts GABA reuptake by astrocytes.

### Histology

#### Tissue preparation, fixation, slice prep, slide mounting

Brains were removed after mice were killed via CO_2_ inhalation and cervical dislocation. For brains, tissue processing for histology began with a 4 hour fixation in 4% paraformaldehyde followed by 3 x 10 min washes in 1x PBS and cryo-protecting the brain in 30% sucrose in PBS overnight at 4°C. Once cryo-protected, the brains were embedded in 3% agar in PBS. 50-µm-thick slices containing the dLGN were created using a Leica VT1000S vibratome, mounted on Superfrost Plus slides (Fisher), and stored at −20°C until stained.

#### Immunostaining and analysis

The slides were rinsed in PBS followed by blocking and permeabilization for 1 hour in blocking buffer (PBS, 0.5% Triton X-100, 5.5% donkey serum, 5.5% goat serum, pH adjusted to 7.4 using NaOH). The slides were then incubated in primary antibody diluted in blocking buffer (1:1000 rabbit-anti-Ankyrin-G) for 3 nights at 4°C. Following incubation with primary antibodies, the slides were washed 6 x 10 min in PBS, incubated in blocking buffer for 1 hour, and incubated with secondary antibody diluted with blocking buffer (1:200 donkey anti-rabbit IgG, Alexa Fluor 568, Invitrogen A-10042). The slides were then washed 3 x 10 min in PBS and 1 x 1 min in dH_2_O, coverslipped with Vectashield Hardset mounting medium, and stored at 4°C until imaged. Ankyrin-G was imaged on a Scientifica two-photon microscope with a MaiTai HP Ti-Sapphire laser tuned to 800 nm with a 123.33 x 123.33 μm field of view (8.303 pixels/μm) centered on the dLGN core. Four images per plane were acquired with 0.5 μm spacing between each plane (laser power at approximately 92 μW).

Images were analyzed using ImageJ, where the images in each plane were grouped and averaged using the “grouped Z project” command with a projection method of “average intensity”. The “brightness/contrast” command was then applied with the “auto” setting to the images. The Simple Neurite Tracer plugin was then used to trace the axon initial segments in each image. The “straighten” command was then applied to each trace, and the intensity profile was plotted for each trace using the “plot profile” command.

#### Optic nerve histology

2-4 mm of optic nerve proximal to the globe was fixed in 4% paraformaldehyde for 3 hours, washed in PBS, and stained with a solution of 2% osmium tetroxide for two hours. Nerves were embedded in paraffin and cross sections obtained at 2 μm thickness by the UNMC Tissue Science Facility. Cross sections were mounted on slides, stained with toluidine blue, and coverslipped with VectaShield Hardset. Whole cross sections of nerves were imaged on an Olympus BX51 WI microscope with a 10x objective (1.68 px/μm). For analyzing glial scarring of the optic nerves ^26^, a region of interest was drawn around the nerve in ImageJ, local contrast was enhanced, and the image was sharpened. The image was thresholded to differentiate darkly-stained regions containing myelinated axons from unstained glial scar. The area of a binary mask comprised of the glial scar was measured and used to calculate the fractional glial scar area of the nerve cross section.

### Statistical Analysis

Statistical analysis was performed using GraphPad Prism 10. A D’Agostino-Pearson test was used to test the normality of data. If non-normally distributed, data were log-transformed prior to analysis or non-parametric approaches were used, as described in the Results. For IOP measurements, a Mann-Whitney test was used to determine significance. A t-test was used to test for statistical significance of histological (glial scarring and axon initial segment length) and pERG data and for sex differences. For experiments involving multiple measurements from individual animals (i.e. patch clamp recordings from multiple cells and axon initial segment measurements), a nested t-test was used to avoid pitfalls from pseudoreplication ^94^. Individual data points (cells, mice, eyes, nerves) along with mean±SEM (standard error of the mean), or median±IQR (inter-quartile range), as indicated in the figure legends.

