## Supplemental figures S1-S5 for "Compensatory responses to glaucoma pathology in the dorsolateral geniculate nucleus"

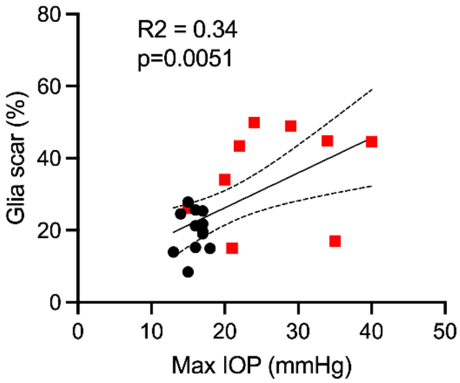


**Supplemental Figure S1 (related to Figure 1) – Correlation of optic nerve glial scarring with peak intraocular pressure in D2 mice.** The extent of glial scarring weakly but significantly correlated with the maximal IOP ($R^2$ = 0.34, p=0.0051), indicative of a relationship between eye pressure and optic nerve injury.

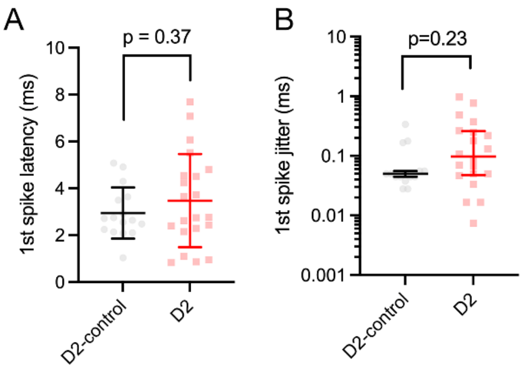


**Supplemental Figure S2 (related to Figure 2) – Latency and jitter of the first action potential evoked in dLGN TC neurons in response to optic tract stimulation.** A) The mean first spike latency, measured as the time from the initiation of the presynaptic stimulus to the initiation of the first action potential did not significantly differ between D2 and D2-control recordings (p=0.37 unpaired nested t-test), although there was a significant increase in cell-to-cell variance (p=0.02, F-test). B) First spike jitter was measured as the absolute value of the difference in time of the first spike from the average time of the first spike for 3-15 repetitions of the stimulus sequence. The first spike jitter for D2 recordings did not significantly differ from that in D2-control recordings (p=0.23, nested t-test of log-transformed values), although the variance was higher in D2 recordings (p=0.024, F-test).

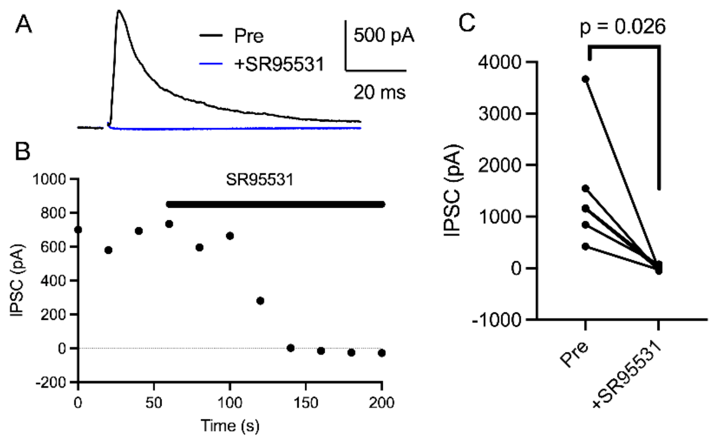


**Supplemental Figure S3 (related to Figure 5) – SR95531 block of feed-forward IPSCs recorded in response to optic tract stimulation.** dLGN TC neurons were voltage-clamped at 0 mV and stimulation delivered to the optic tract. A) example traces before and after SR95531 (25 μM) application. B) Time course an experiment showing amplitudes of IPSCs evoked at 20-second intervals before and during bath application of SR95531. C) Group data. The IPSCs were entirely blocked by 25 μM SR95531 (p=0.026, paired t-test).

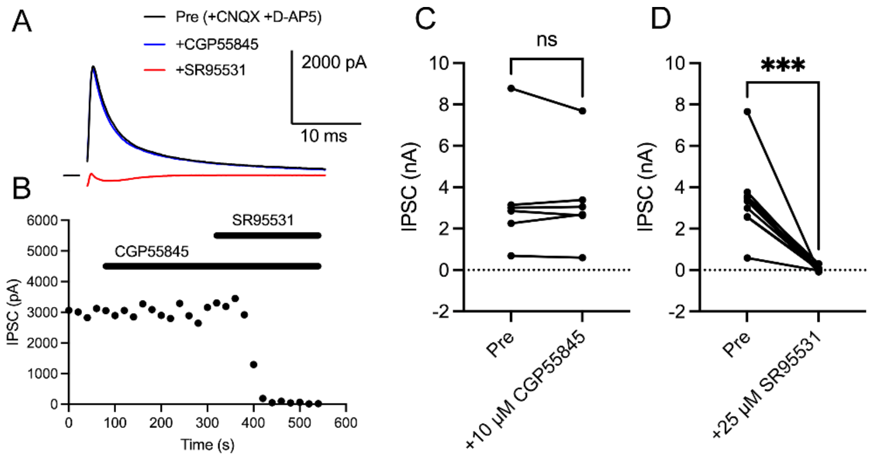


**Supplemental Figure S4 (related to Figure 5) – SR95531 block of IPSCs evoked by extracellular stimulation without involvement of $GABA_B$ receptor signaling.** A) IPSCs recorded in a TC neuron ($V_{hold}$ = 0 mV) evoked by extracellular stimulation with an aCSF-filled patch pipette located ~25 microns from the cell body in the presence of CNQX (20 μM) and D-AP5 (50 μM). Traces show baseline, after application of the $GABA_B$ receptor antagonist CGP55845 (10 μM) and after application of SR95531 (25 μM). B) Time course of an experiment showing IPSC amplitude measurements in baseline and after addition of CGP55845 and SR95531. C) CGP55845 had no significant effect on IPSC amplitude (n=6; p=0.62, paired t-test). D) IPSCs were entirely blocked by SR95531 (n=10; p=0.00020, paired t-test).

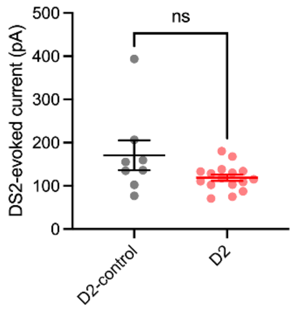


**Supplemental Figure S5 (related to Figure 6) – DS2-evoked outward current.** Measurement of the change in baseline holding current evoked by DS2 application (10 μM) while TC neurons were voltage-clamped at the cationic reversal potential ($V_{hold}$ = 0 mV) and in the presence of CNQX (20 μM) and D-AP5 (50 μM). DS2 bath application led to an outward current in recorded dLGN TC neurons. Although the amplitude of the DS2-evoked current was slightly larger in D2-control TC neurons compared to those from D2 mice, the difference was not significant (p=0.061, nested t-test).
